# Sfrp1 deficiency makes retinal photoreceptors prone to degeneration

**DOI:** 10.1101/831479

**Authors:** Elsa Cisneros, Fabiana di Marco, Javier Rueda-Carrasco, Concepción Lillo, Guadalupe Pereyra, María Jesús Martín-Bermejo, Alba Vargas, Rocío Sanchez, África Sandonís, Pilar Esteve, Paola Bovolenta

**Affiliations:** Centro de Biología Molecular Severo Ochoa, CSIC-UAM, Madrid, Spain; Centro de Investigación Biomédica en Red de Enfermedades Raras (CIBERER). Madrid, Spain; Departamento de Biología Celular y Patología. Universidad de Salamanca, Instituto de Neurociencias de Castilla y León and IBSAL. Salamanca, Spain

**Keywords:** Retina, Outer Limiting Membrane, Cell adhesion, Photoreceptor degeneration, Light-induced damage, Neurodegenerative diseases, Aging

## Abstract

Millions of individuals worldwide suffer from impaired vision, a condition with multiple origins that often impinge upon the light sensing cells of the retina, the photoreceptors, affecting their integrity. The molecular components contributing to this integrity are however not yet fully understood. Here we have asked whether Secreted Frizzled Related Protein 1 (SFRP1) may be one of such factors. SFRP1 has a context-dependent function as modulator of Wnt signalling or of the proteolytic activity of A Disintegrin And Metalloproteases (ADAM) 10, a main regulator of neural cell-cell communication. We report that in *Sfrp1^-/-^* mice, the outer limiting membrane (OLM) is discontinuous and the photoreceptors disorganized and more prone to light-induced damage. *Sfrp1* loss significantly enhances the effect of the *Rpe65^Leu450Leu^* genetic variant -present in the mouse genetic background- which confers sensitivity to light-induced stress. These alterations worsen with age, affect visual function and are associated to an increased proteolysis of Protocadherin 21 (PCDH21), localized at the photoreceptor outer segment, and N-cadherin, an OLM component. We thus propose that SFRP1 contributes to photoreceptor fitness with a mechanism that involves the maintenance of OLM integrity. These conclusions are discussed in view of the broader implication of SFRP1 in neurodegeneration and aging.

## Introduction

Photoreceptors are specialized neurons localized in the outermost layer of the retina, which are responsible for converting light information into electrical activity. There are two types of photoreceptors: cones that mediate vision in bright light (photopic vision), and rods that support vision in dim light (scotopic vision). Each photoreceptor develops a highly modified cilium, the outer segment (OS), composed of a stack of membranes that contains light-sensing visual pigments and other photo-transduction proteins connected to the photoreceptor cell body by the inner segment (IS). The organization, polarity and function of these specialized structures is assured at the proximal side by the presence of the outer limiting membrane (OLM), composed of a series of adherens junctions between the initial portion of the IS and the neighbouring Müller glial end-feet. At the distal side, instead, the OSs are in contact with the retinal pigment epithelium (RPE), a monolayer of cells that daily phagocyte the OS abutting portion and provides metabolic support to the photoreceptors ^1^. Thus, the physiology of photoreceptors is critically dependent, among others, on their interaction with adjacent non-neuronal cell types. Environmental, age-related and genetic defects disrupting these interactions lead to the progressive death of photoreceptors and thereby to the progressive loss of sight, often followed by total blindness ^2^. Indeed, loss of OLM integrity is commonly observed in different forms of retinal dystrophies, including Retinitis Pigmentosa (RP) ^2^, a collection of monogenic disorders characterized by a tremendous genetic heterogeneity ^3^. Furthermore, RP causative genes include those coding proteins enriched at the OS and OLM, such as PCDH21 or Crumbs homolog 1 (CRB1) ^4–6^ and even metalloproteases, such as ADAM9 ^7^, known to cleave different cell adhesion molecules ^8,9^. The molecular components that maintain photoreceptors’ interaction with the nearby tissues, thereby contributing to their integrity, are however not yet fully understood. The identification of such mechanisms may be instrumental in designing strategies for slowing down the progression of photoreceptors’ degeneration and perhaps explain the observed inter-individual variability of photoreceptor degeneration even among relatives carrying identical mutations or risk alleles ^10^ in the case of genetic disorders.

We have recently shown that elevated levels of Secreted Frizzled-Related Protein (SFRP) 1-a secreted protein with the dual function of modulating Wnt signalling ^11^ and regulating the enzymatic activity of the α-sheddase ADAM10 ^12^-contributes to the pathogenesis of Alzheimer’s Disease ^13^. Consistent with the notion that ADAM10 cleaves a large number of substrates -including the amyloid precursor protein (APP) and proteins involved in neuroinflammation and synaptic plasticity ^14, 15^-, high levels of SFRP1 prevent ADAM10-mediated non-amyloidogenic processing of APP ^13^ and favour neuroinflammation ^16^. *Sfrp1* and its homologues *Sfrp2* and *Sfrp5* have been implicated in different aspects of eye and retinal development ^17–23^. Furthermore, SFRP1 has been found to be expressed in the adult retina, mostly localized to the photoreceptor layer ^25^, in contrast to what observed in the brain, in which SFRP1 expression is minimal in homeostatic conditions ^13, 24^. *SFRPs* genes are an unlikely primary cause of RP in humans ^26^; however, their expression is notably increased and ectopically distributed in the retinas of patients with RP ^25, 27^. These observations, together with the notion that variation of *SFRP* expression has been noted in a variety of other pathological conditions ^28^, made us ask whether SFRP1 may be involved in maintaining photoreceptors’ integrity.

Here we report here that SFRP1 helps maintaining photoreceptor’s integrity. In its absence, photoreceptors of young and mature mice show subtle morphological alterations of the OS associated with discontinuities of the OLM and an increased proteolytical processing of two of its components: N-cadherin and PCDH21. Furthermore, *Sfrp1* absence significantly increases the sensitivity of the photoreceptors to light-induced damage in the presence of the sensitizing *Rpe65^Leu450Leu^* gene variant, present in the genetic background of the mice.

## Results

### Young adult Sfrp1^-/-^ retinas show subtle defects in cone photoreceptor organization

The final goal of our study was to determine whether SFRP1 is part of the molecular machinery that maintains photoreceptors’ integrity. We reasoned that, if this is the case, mice lacking SFRP1 activity (*Sfrp1^-/-^*) should have a different susceptibility than their wild type (wt) counterparts to light-induced damage, taken as a frequently used experimental paradigm to study photoreceptor degeneration ^29^. *Sfrp1* does not seem to be required for mouse retinal development ^18, 30, 31^. However, there are somewhat controversial reports on its adult retinal localization ^32, 33^ and, to our knowledge, no specific information on its possible function in adult retinal homeostasis. We addressed these issues first.

To clarify *Sfrp1* distribution, we hybridized sections of 1 month-old mouse eyes with a specific probe. Our results supported the report by Liu et al. ^32^. High levels of *Sfrp1* were localized to the inner nuclear (INL) and ganglion cell (GCL) layers. Lower levels were also found in the outer nuclear (or photoreceptor) layer (ONL; Fig. 1a), with a more abundant distribution in a subset of cells located at the outermost region (Fig. 1b, white arrows), in which the cell bodies of cone photoreceptors are found ^34^. Immunostaining with specific antibodies confirmed a similar layered distribution of the protein that, according to its secreted and dispersible nature ^18, 35^, was notably accumulated at the OLM/OS region (Fig. 1c,d). ELISA determination of SFRP1 content in retinal extracts from mice of age comprised between 1 and 25 months showed that its levels significantly decreased with age (Fig. 1k). *In situ* hybridization analysis did not detect the expression of the closely related *Sfrp2* in the adult mouse retina as described for humans ^27^, whereas *Sfrp5* reporter expression was localized only in few sparse retinal cells and in the retinal pigmented epithelium (Fig. S1).

**Figure 1.**
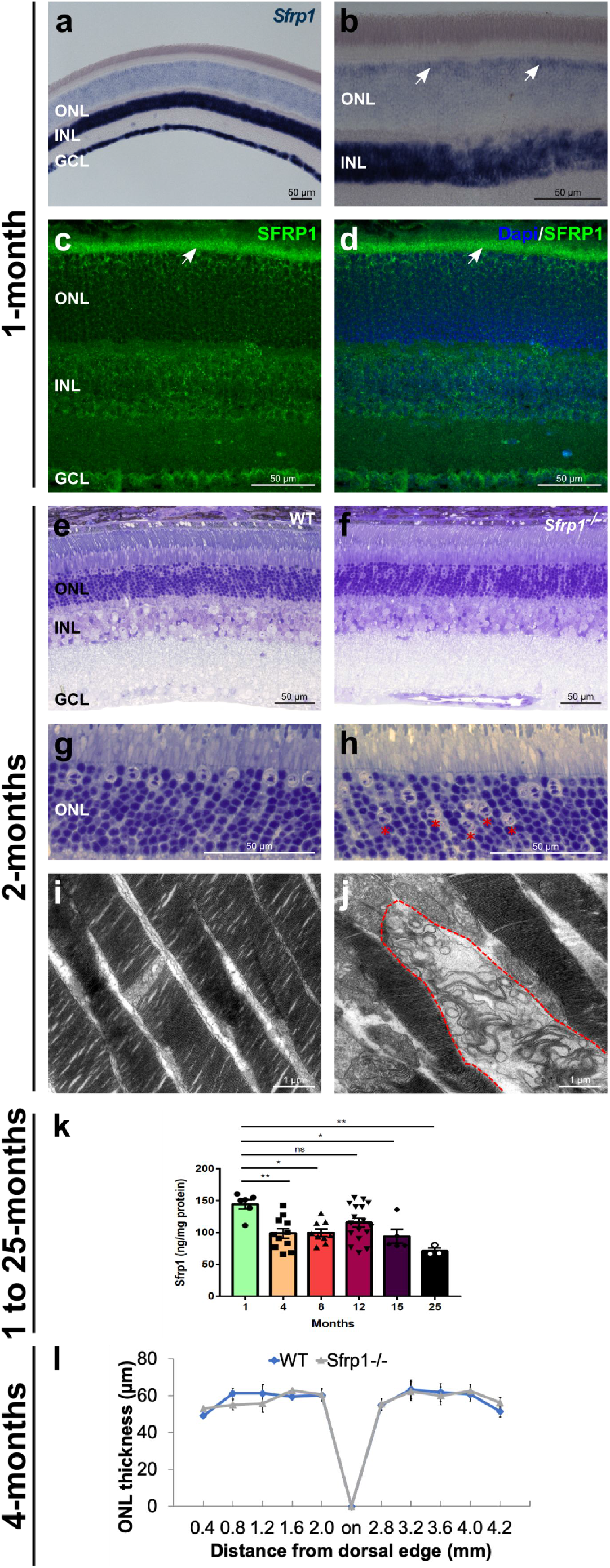
SFRP1 is expressed in the retina and required for photoreceptor fitness. **a-d)** Frontal cryostat sections from 1 month-old wt animals hybridized (a,b), or immunostained (c,d) for SFRP1 and counterstained with DAPI (d). The white arrows in b) indicate the increased signal in the outermost region of the ONL. Note Sfrp1 accumulation in the OLM (arrowheads in c,d). **e-h)** Semi-thin frontal sections from 2 months-old wt and *Sfrp1^-/-^* retinas stained with toluidine blue. Note the abnormal localization of cone nuclei in the mutant retina (red asterisks in h). **i, j)** TEM analysis of the OS. Note that in the mutants, but not in wt, the stacks of membranes of the OS are disorganized (area enclosed in dotted line in j). **k)** ELISA analysis of SFRP1 levels in total protein extracts from 1 to 25 months-old retinas from wt mice. Note the significant decrease of SFRP1 content as the animals age; One-way ANOVA, with post hoc Bonferroni analysis; *: p<0.05, **: p<0.01. **l)** The graph shows the ONL thickness in 4 months-old wt and *Sfrp1^-/-^* retinas (measures were taken in cryostat frontal sections of the eye at the level of the optic disk). No significant differences were detected between wt and *Sfrp1^-/-^* retinas (Mann-Withney U test). Abbreviations: GCL: ganglion cell layer; INL: inner nuclear layer; on: optic nerve; ONL: outer nuclear layer.

To determine if the activity of SFRP1 is required for the organization of the adult retina, we compared the retinal morphology of 2 months-old *Sfrp1^-/-^* and wt mice using toluidine blue stained semi-thin sections. No appreciable gross differences were detected between the two genotypes and the thickness of the ONL layer was comparable in both the dorsal and ventral quadrants (Fig. 1e,f), even in 4 months-old mice (Fig. 1l; n=3 mice / genotype, Mann-Whitney U test). In mice, cone cell bodies are mostly arranged in a single row below the OLM with nuclei characterized by the presence of 1-3 irregular clumps of heterochromatin. Rods instead occupy all the ONL rows with more densely stained nuclei and a single large central clump of heterochromatin ^34^. This arrangement was easily recognizable in wt retinas observed at higher magnification (Fig. 1g), with 86.1% (± 0.2) of the cones localized to the first or second row of the ONL, whereas the remaining 13.9% (± 2.9) were found in other rows (n=3 mice). In *Sfrp1^-/-^* retinas instead, cone nuclei were abnormally distributed (Fig. 1h, red asterisks) and only 51.1% (± 6.2) of the cone cell bodies were found in the first two rows. The remaining 48.9% (± 3.1) were distributed in other rows, with a significant cone increase in deeper rows (n=3 mice; Chi-square test, p<0.001). Nonetheless, the total number of cones was not significantly different between wt (291 ± 8, in nerve head sections, n=3) and *Sfrp1^-/-^* mice (336 ± 13.5, n=3; Mann-Whitney U test), at least in young adults. Nevertheless, ultra-structural analysis revealed occasional degenerative signs in the OS of the mutants, which were never observed in wt (Fig. 1i,j).

*Sfrp1^-/-^* mice were bred in a mixed C57BL/6 x 129 background. Previous studies have warned that C57BL/6N sub-strain carries a single nucleotide deletion in the Crumb homologs 1 (*Crb1)* gene, which causes a retinal phenotype that may override that of other genes of interest ^36^. Our animals were bread in the C57BL/6J sub-strain, reported to carry a wt allele ^36^. Nevertheless, and to confirm that the observed defects were bona fide associated to the loss of *Sfrp1* function, we sequenced the *Crb1* gene in the *Sfrp1^-/-^* line. No *Crb1* mutations were found in all the analysed samples of our colony (not shown).

Once excluded the possible contribution of a defective *Crb1* allele, we next asked if the subtle defects observed in *Sfrp1^-/-^* retinas had an impact on photo-transduction or synaptic transmission between photoreceptors and bipolar cells. To this end, we compared the electrical responses of groups of 4-months old wt and *Sfrp1^-/-^* mice (n=12 per genotype) to light stimuli of increasing intensity and under increasing frequency. ERGs were recorded under scotopic (Fig. 2a-d) and photopic (Fig. 2e,f,g) conditions to evaluate rod/rod bipolar cells and cone/cone bipolar function, respectively. Mutant mice showed a small but statistically non-significant (Mann-Whitney U test) decrease of photoreceptor (a-wave, see Fig S2) scotopic (Fig. 2c,d) and photopic (Fig. 1f) response at higher intensities and a tendency towards a lower response under flicker stimuli of low frequency (Fig. 2g).

**Figure 2.**
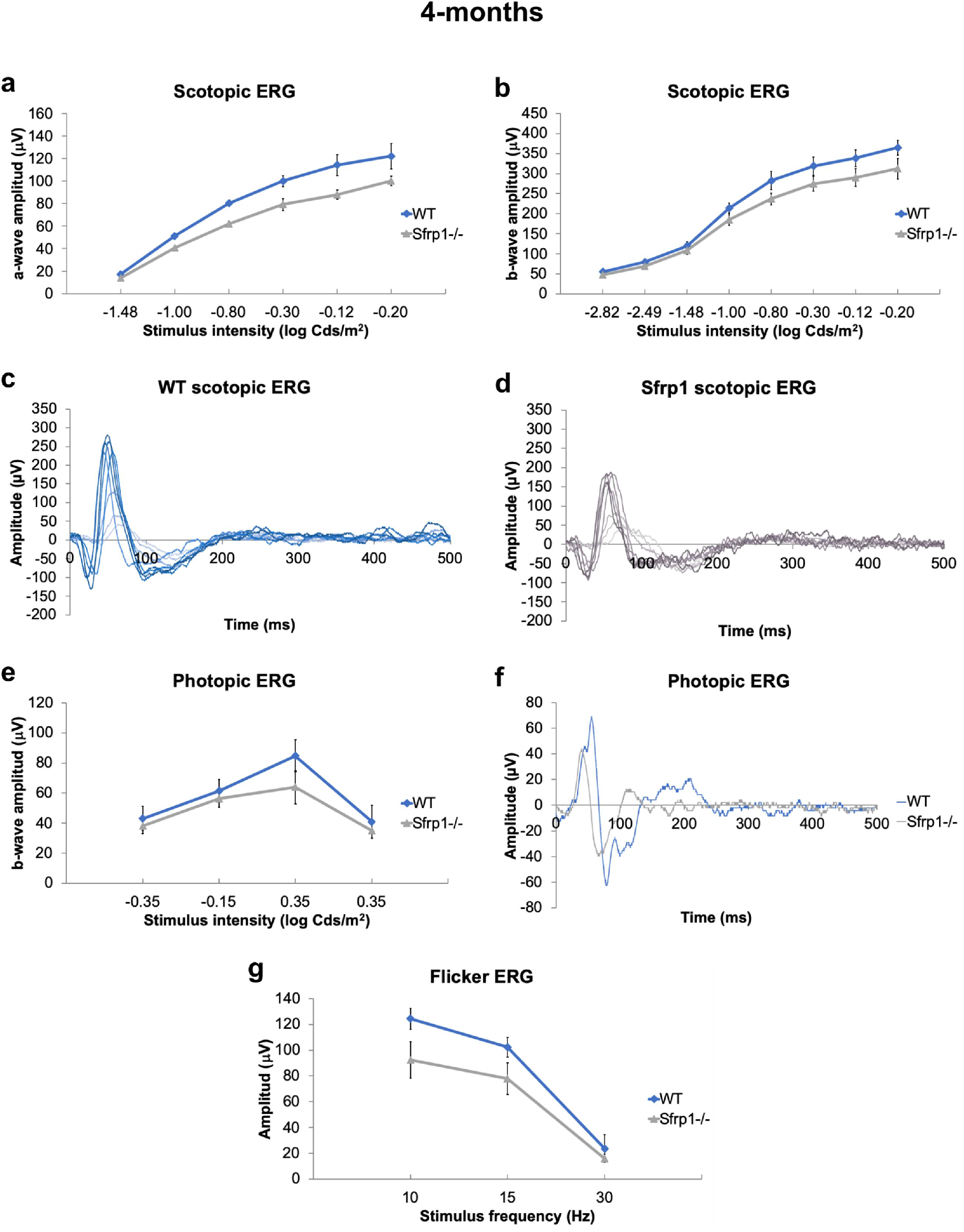
*Sfrp1^-/-^* mice have a slightly reduced visual function. **a-d**) The graphs and traces of representative recordings show the a- and b-wave amplitude (μV; defined in Fig S2) of the scotopic ERG (rods, a-d) and the b-wave amplitude of photopic ERG (cones, e,f) from 4 months-old wt and *Sfrp1^-/-^* animals to flash light stimuli of increasing brightness (cd/m^2^). **e**) The graph shows the amplitude response (μV) to flicker stimuli of increasing frequencies (Hz). No significant changes were detected (Mann-Whitney U test) but mutant mice show a reduced response.

Together these data suggest that SFRP1 contributes to maintain retinal organization but its absence has negligible effects on young adults.

### Retinal disorganization and visual performance of Sfrp1^-/-^ retinas worsen with age

Photoreceptors disorganization, decreased OS length and scotopic responses together with reduced synaptic density and loss of melanin granules are among the alterations found in the aging mammalian retinas ^37–39^. The retinal phenotype of the young adult *Sfrp1^-/-^* mice shared some of these features, suggesting that the observed features could worsen with time.

To address this possibility gross retinal morphology of 6 months-old wt and *Sfrp1^-/-^* mice was analysed, finding no considerable macroscopic differences. However, the well-aligned columnar organization of the photoreceptors observed in wt was lost in the *Sfrp1^-/-^* retinas and, cone photoreceptor nuclei were distributed throughout the width of the ONL (Fig. 3a,b), as we observed in young mice. Many photoreceptors’ nuclei were also misplaced past the OLM that normally overlays the ONL (n=6, Fig. 3c,d). Retinal immunostaining with the Trasducin/GNB3 and rhodopsin (Rho), markers of the OS of cones and rods, respectively, confirmed subtle photoreceptors’ alterations. Trasducin/GNB3 staining was particularly reduced and fragmented in the mutant retinas (n=3 mice for each age, Fig. 3e,f), reflecting a small but significant reduction of the number cone nuclei counted in toluidine blue stained sections (wt, 231.6 ± 12 vs. *Sfrp1^-/-^* 149 ± 26.7, in retinal nerve head sections; n=3 and 5 mice, Mann Whitney U test, p=0.025). In wt mice, Rho normally localises to the OS (Fig. 3g), but in the *Sfrp1^-/-^* retinas labelling was also observed in the cell bodies of a number of cells (Fig. 3h). Although ONL cell density was an obstacle for precise evaluation of the proportion of affected rods, cells with abnormal Rho distribution were consistently observed in both eyes of all the analysed animals (n=3 mice for each age analysed, i.e., 6 and 8 month).

**Figure 3.**
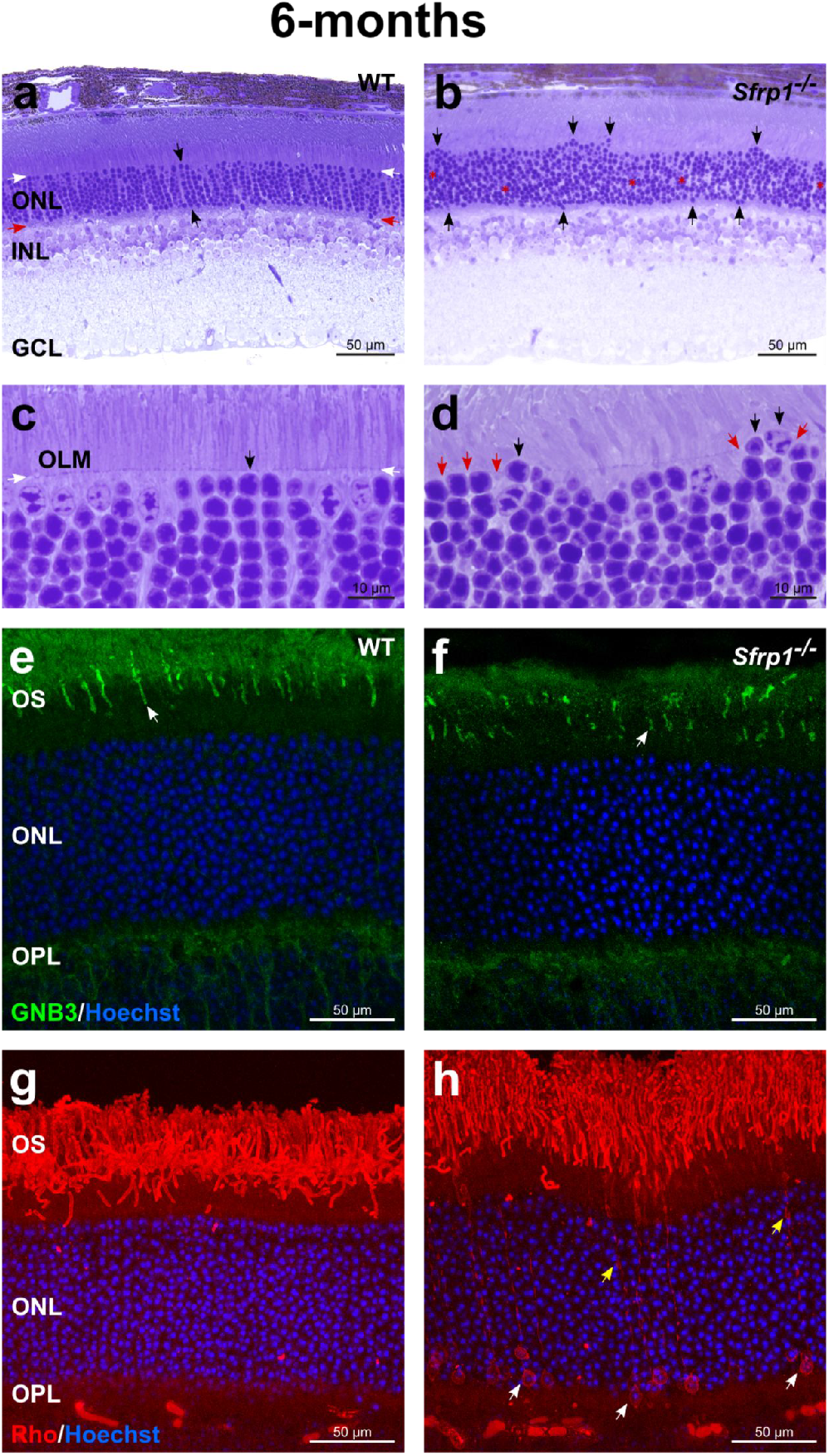
Morphological alterations of *Sfrp1^-/-^* retinas become more evident with age. **ad)** Semi-thin plastic frontal sections from 6 months-old wt and *Sfrp1^-/-^* retinas stained with toluidine blue. Black arrows in a,c indicate an organized column of photoreceptors, whereas white and red arrows indicate the outer and inner limit of the ONL. In b and d black and red arrows point to misplaced photoreceptor nuclei and discontinuous OLM respectively. **e-h)** Frontal cryostat sections from 6 months-old wt and *Sfrp1^-/^*^-^ retinas immunostained with anti GNB3 (cones) or Rho (rods) antibodies and counterstained with Hoechst. White arrows in e,f point to the OS. Yellow and white arrows in h indicate accumulation of Rho in rod processes and cell bodies respectively. Abbreviations. GCL: ganglion cell layer; INL: inner nuclear layer; on: optic nerve; ONL: outer nuclear layer.

The presence of Rho in the cell bodies reflects cell intrinsic or OLM disturbances ^40^. Indeed, the OLM seals off the photoreceptor inner and outer segments from the rest of the cell, thereby limiting the diffusion of the photo-transduction cascade components ^41^. This notion together with the displacement of the photoreceptor nuclei (Fig. 3c, d), similar to that observed upon pharmacological disruption of the OLM ^42, 43^, suggested the existence of OLM alterations in the mutants. The intermittent brakeage of the OLM already apparent in histological sections (red arrows in Fig. 3d) was confirmed by ultrastructural analysis (n=3). The well-organized arrangement of zonula adherens junctional complexes between the plasma membranes of the photoreceptor ISs and the apical processes of Müller glia that compose the OLM was readily visible in wt retinas (Fig. 4a). This arrangement was no longer present in those regions of the *Sfrp1^-/-^* retinas in which photoreceptors’ nuclei were displaced bearing signs of degeneration in both the cell body (Fig. 4b) and OS. Defective outer segments were more abundant than those observed at 2 months of age (Fig. 1j) and mostly localized in the central retina as determined by quantification in semi-thin sections (Fig. 4c, d; 5 ± 1 in wt and 25 ± 3 in *Sfrp1^-/-^* nerve head sections, n=3, Mann-Whitney U test, p=0.043). In *Sfrp1^-/-^* retinas, RPE cells presented less and smaller melanosomes (Fig 4e,f) and synaptic contacts between photoreceptors and bipolar interneurons were also altered, with an abnormal accumulation of synaptic vesicles (Fig. 4h) and the degeneration of pre and post-synaptic terminals (red arrows in Fig. 4i,l). None of these defects were observed in wt retinas (Fig. 4g) but have been reported in aging retinas ^37–39^.

**Figure 4.**
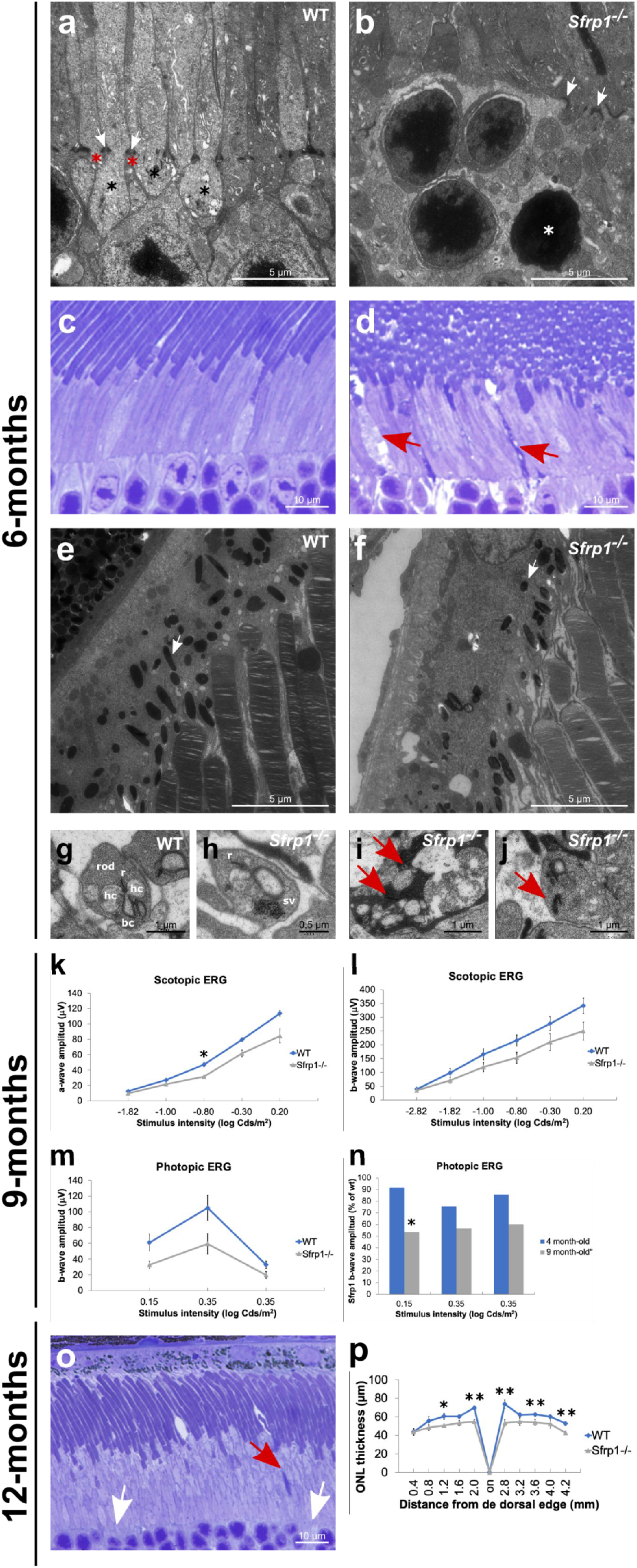
OLM is disrupted in the retina of *Sfrp1^-/-^* mice in association with an impoverishment of visual function. **a,b; e-j)** TEM analysis of the retina from 6 months-old wt and *Sfrp1^-/-^* mice. Images show the OLM (a,b), RPE (e,f) and photoreceptor-bipolar cells synapses (g-j). White arrows in a,b point to adherens junctions between Müller glial cell endfeet (red asterisks) and photoreceptors (black asterisks). White arrows in e,f point to melanosomes. Red arrows in i,j indicate synaptic terminal disorganization. **c-d)** Semi-thin frontal sections from 6 months-old wt and *Sfrp1^-/-^* retinas stained with toluidine blue. Note the abnormal presence of degenerating outer segments in the mutant retina (red arrows in d). **k-l**) The graphs show the ERG recordings of rods and rod bipolar cells (scotopic, k,l) and cone - pathway (photopic, m) response to light of 9 months-old wt and *Sfrp1^-/-^* animals. **n)** The graph represents the activity of photoreceptors from 4 and 9 months-old *Sfrp1^-/-^* mice. Note that activity is expressed as a percentage of wt age-matched animals. Only 9 months-old Sfrp1 mice showed a significantly reduced activity relative to their control. **o)** Semi-thin frontal sections from 10 months-old *Sfrp1^-/-^* retina stained with toluidine blue. Note the continuous presence of degenerating (red arrows) and enlarged OS with evident breakage of the OLM (white arrows); **p)** The graphs represent the thickness of the ONL in 12 months-old wt and *Sfrp1^-/-^* retinas showing a significant reduction in the mutants more evident in central regions. Mann-Whitney U test; * p<0.05, ** p<0.01. Abbreviations. bc: bipolar cell; hc: horizontal cell; r ribbon; sv: synaptic vesicles.

Recording of ERGs responses (Fig S2) under scotopic (Fig. 4k,l) and photopic (Fig. 4m,n) conditions in 9 months-old wt and *Sfrp1^-/-^* mice showed that the above described morphological defects of *Sfrp1^-/-^* mice were associated with a poorer performance of rods as compared to wt mice (Fig. 4k; n=5 per genotype, Mann-Whitney U test, p=0.035). Direct comparison of the photopic responses of 4 and 9 months-old *Sfrp1^-/-^* mice indicated a progressive deterioration with aging (Fig. 4n; Mann-Whitney U test, p=0.033).

Morphological analysis of 12 months-old mice showed worsening of photoreceptor damage with enlarged and degenerated OS and an evident breakage of the OLM (Fig. 4o). Overall, the thickness of ONL was significantly reduced (20% on average) across the almost entire dorso-ventral axis of the retinas as compared to age-matched wt (Fig. 4r; n=5 per genotype, Mann-Whitney U test, p<0.05).

Together these data indicate that *Sfrp1* loss causes a slow but progressive deterioration of the retinal integrity associated with decrease of visual function.

### The proteolysis of OLM and OS components is increased in Sfrp1^-/-^ retinas

The region of the OLM is enriched in cell adhesion molecules, such as N-cadherin found at the adherens junctions ^44^ and protocadherin PCDH21, located at the base of the OS ^45^. Other transmembrane molecules are, for example, CRB2 and CRB1, found at the subapical region (right above the adherens junctions) of the photoreceptors and/or Müller glial cells as part of a large protein complex ^6, 46^. As already mentioned, a tight adhesion between IS of the photoreceptors and the end feet of the Müller cells is critical for photoreceptor integrity and retinal organization ^47^. Furthermore, variation of the human CRB1 and CRB2 genes are responsible of retinal dystrophies such as RP ^48^ and inactivation of their murine homologues disrupts the OLM and causes photoreceptor loss ^49^. We thus reasoned that the retinal phenotype of the adult *Sfrp1^-/-^* mice could be mechanistically linked to poor Müller/photoreceptor cell adhesion, which, in turn, would be responsible of photoreceptor alterations. This mechanism is based on the notion that SFRP1 acts as a negative modulator of ADAM10 ^12, 13, 22, 24^, which proteolytically removes the ectodomains (ectodomain shedding) of a large number membrane proteins ^8^, including cell adhesion molecules such as Cadherins and proto-Cadherins ^14, 50, 51^, thereby controlling cell-cell interactions ^8, 51^. Consistent with this idea, it has been shown that abnormal proteolytical processing of OS components influences photoreceptor degeneration ^52^. Furthermore, different ADAM proteins, including ADAM10, have been found expressed in different retinal layers and cell types, comprising Müller cells ^53, 54^. We confirmed this distribution with immunohistochemical analysis of ADAM10 localization. In 3 months-old retinas ADAM10 was specifically found in ganglion cells, in the INL (presumably in Müller glial cells based on position) and in the OLM (Fig 5a,b). Thus and given that SFRP1 is particularly abundant in the OLM/OS region (Fig. 1c,d), we hypothesized that, in SFRP1 absence, component of the OLM could be proteolyzed more efficiently.

We therefore tested if an enhanced proteolytical processing of N-cadherin and/or PCDH21 could explain the retinal alterations of *Sfrp1^-/-^* mice. To this end, we analysed using Western blot the proportion of proteolyzed versus full-length protein levels of N-cadherin and PCDH21 in wt, *Sfrp1^-/-^* and *rd10* mice. These mice, carrying a missense mutation in the rod specific *Pde6b*, are a widely used model of fast photoreceptor degeneration ^55^ and thus appeared useful to assess mechanistic specificity. Proteolytical processing of full-length N-cadherin (135 kD) results in the generation of a C-terminal intracellular peptide of 35 kDa ^51^, whereas the 120 kDa full length PCDH21 is cleaved in two fragments of 95 (N-terminal) and 25 (C-terminal) kDa ^52^. Antibodies against the C-terminal portion of either N-Cadherin or PCDH21 were probed against retinal extracts of 3 weeks-old mice. At this stage, the majority of photoreceptors in *rd10* mice are still alive, allowing for comparison. *Sfrp1^-/-^* retinal extracts contained an increased proportion of the C-terminal fragments of both N-cadherin and PCDH21 when compared to those detected in wt (Fig. 5c,d; Fig. S3, about 2- and 3.5-fold, respectively). N-cadherin but not PCDH21 proteolysis was also observed in *rd10* retinas (Fig. 5c,d; Fig. S3). Comparable and unprocessed levels of CRB1 were present in the 3 genotypes (Fig. 5e; Fig. S3), supporting processing specificity. Furthermore, the signal of immunostaining for N-cadherin and β-catenin -which functionally interacts with cadherins at the adherens junctions ^56^-appeared discontinuous and spread along the Müller glial processes in the *Sfrp1^-/-^* retinas (Fig. 5g,i) but as a continuous line in wt retinas (Fig. 5f,h). Staining of Müller glial cells with anti-Glutamine Synthetase (GS) confirmed the discontinuous arrangement of their apical end-feet in the mutants but not in the wt retinas (Fig. 5j,k).

**Figure 5.**
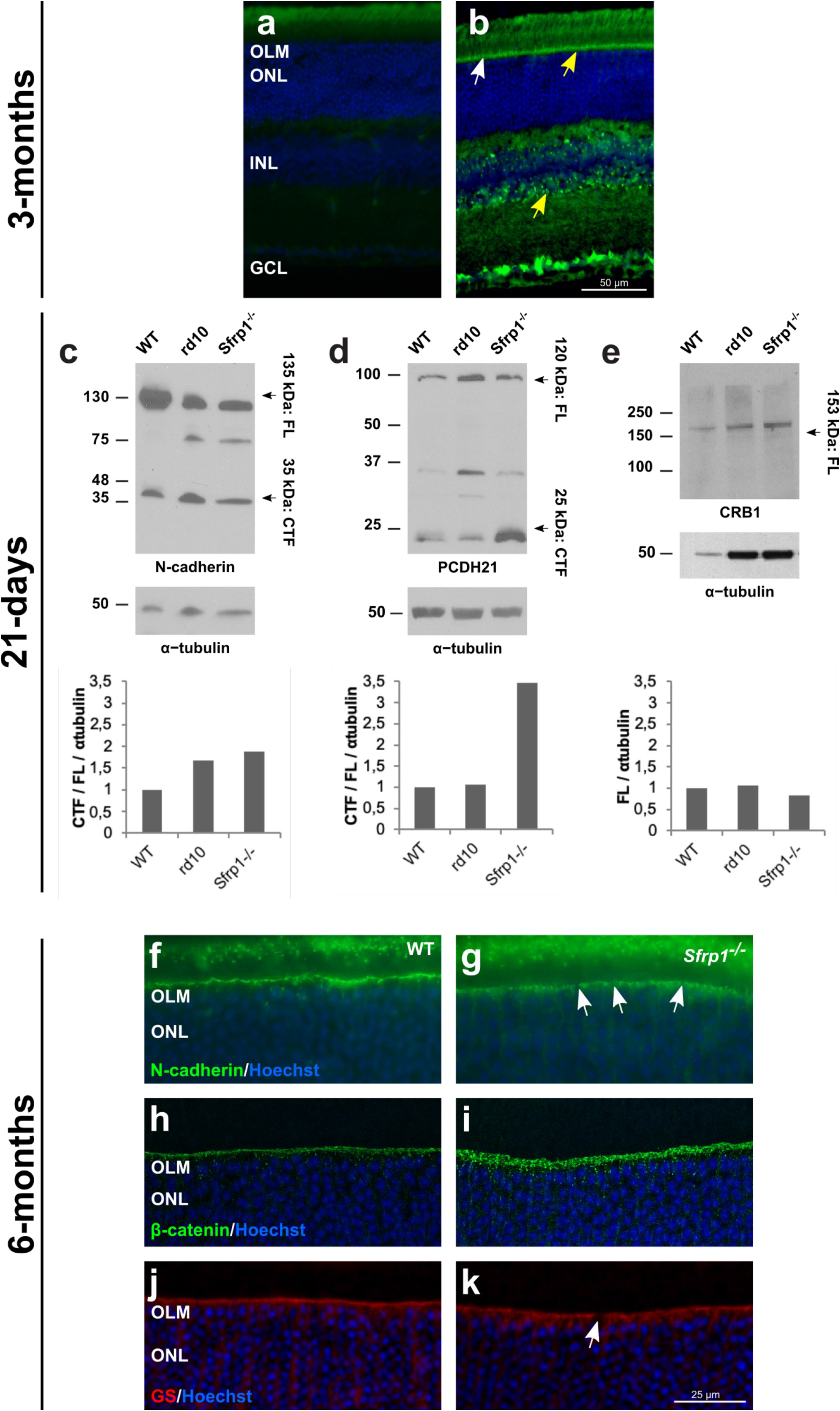
Proteolysis of OLM and OS proteins is increased in *Sfrp1^-/-^* retinas leading to an abnormal protein distribution. **a, b)** Frontal cryostat sections from 3 months-old wt retinas immunostained with secondary antibodies only (a) or with anti-ADAM10 (b). Sections are counterstained with Hoechst. Note the specific immune signal in the OLM (arrows) and in the INL where Müller glial cells are located (yellow arrows). **c-e)** Western blot analysis of N-cadherin, PCDH21 and CRB1 content in total protein extracts from three weeks-old wt, *Sfrp1^-/-^* and *rd10* retinas, as indicated in the panels. For N-cadherin and PCDH21, membranes were probed with antibodies recognizing the respective C-terminal fragments (CTF). Graphs at the bottom of each panel indicate the rate of CTF proteolysis for N-cadherin (c) and PCDH21 (d), calculated as CTF/FL, or the CRB1 levels (e). Plotted values are normalized to α-tubulin. **f-k**) Frontal cryostat sections from 6 months-old wt and *Sfrp1^-/-^* retinas immunostained with antibodies against N-cadherin, β-catenin, and GS as indicated in the panels. Sections are counterstained with Hoechst. White arrows in g, k indicate the gaps in the OLM. Scale bar calibration in k applies to all panels. Abbreviations: CTF: C-terminal fragment; FL: full length; OLM: outer limiting membrane; ONL: outer nuclear layer.

Altogether these data indicate that SFRP1 indirectly regulates photoreceptors/Müller glia interaction, which is, in turn, essential for retinal integrity and optimal function.

### Sfrp1 bestows resistance to the photoreceptors upon light induced damage

If SFRP1 maintains photoreceptor integrity, in its absence photoreceptors should be more sensitive to a neurodegenerative stimulus. Exposure to high intensity light is an experimental paradigm widely used to induce a rhodopsin mediated stereotypic photoreceptor apoptosis ^29^, to which cones are less susceptible ^57, 58^. The degree of response however varies according to the pigmentation and genetic background of the animals ^29^. Most prominently, a Leu450Met variation in the RPE-specific *Rpe65* gene, found in the C57BL/6 and 129 mouse strains ^59^, confers resistance to light damage. Given the genetic background of the *Sfrp1^-/-^* mice, we sequenced the *Rpe65* gene in our colony, finding the presence the *Rpe65^Leu450Met^* variant mostly in heterozygosis. To clearly separate the contribution of the two alleles we generated *Rpe65^Leu450Leu^;Sfrp1^-/-^* and *Rpe65e^Leu450Met^;Sfrp1^-/-^* animals and their respective controls.

We thereafter exposed 4 months-old wt and *Sfrp1^-/-^* mice carrying the two *Rpe65* variants to a 15.000 lux light source for 8 h and analysed the extend of photoreceptors’ loss by immunohistochemical analysis 7 days after treatment. Under these conditions and compared to untreated animals (the two *Rpe65* variant were undistinguishable), the retinas of *Rpe65e^Leu450Met^;Sfrp^+/+^* mice were only slightly affected. The morphology and thickness of their ONL as well as the immunolabelling for rod (Rho) and cones (GNB3) markers were comparable to that of untreated animals (Fig. 6a,c,e,g,m). Given the absence of the resistance allele, the photoreceptors of *Rpe65e^Leu450Leu^;Sfrp^+/+^* mice instead presented signs of disorganization and the ONL was slightly thinner in central regions as compared to untreated mice (Fig. 6a,c,i,k,n). By contrast, the retinas of *Rpe65e^Leu450Met^;Sfrp1^-/-^* and, to a larger extend, those of *Rpe65^Leu450Leu^;Sfrp1^-/-^* mice were visibly affected by light damage. There was a central^high^ to peripheral^low^ loss of both rod and cones photoreceptors with abnormal localisation of Rho in the cell bodies (Fig. 6b,d,f,h,j,l,n). Cone loss appeared proportionally more significant than that of rods in mice carrying the *Rpe65^Leu450Leu^* vs the *Rpe65e^Leu450Met^* variant (compare Fig. 6f,h with j,l; Kruskal Wallis with post hoc Dunn, p<0.05).

**Figure 6.**
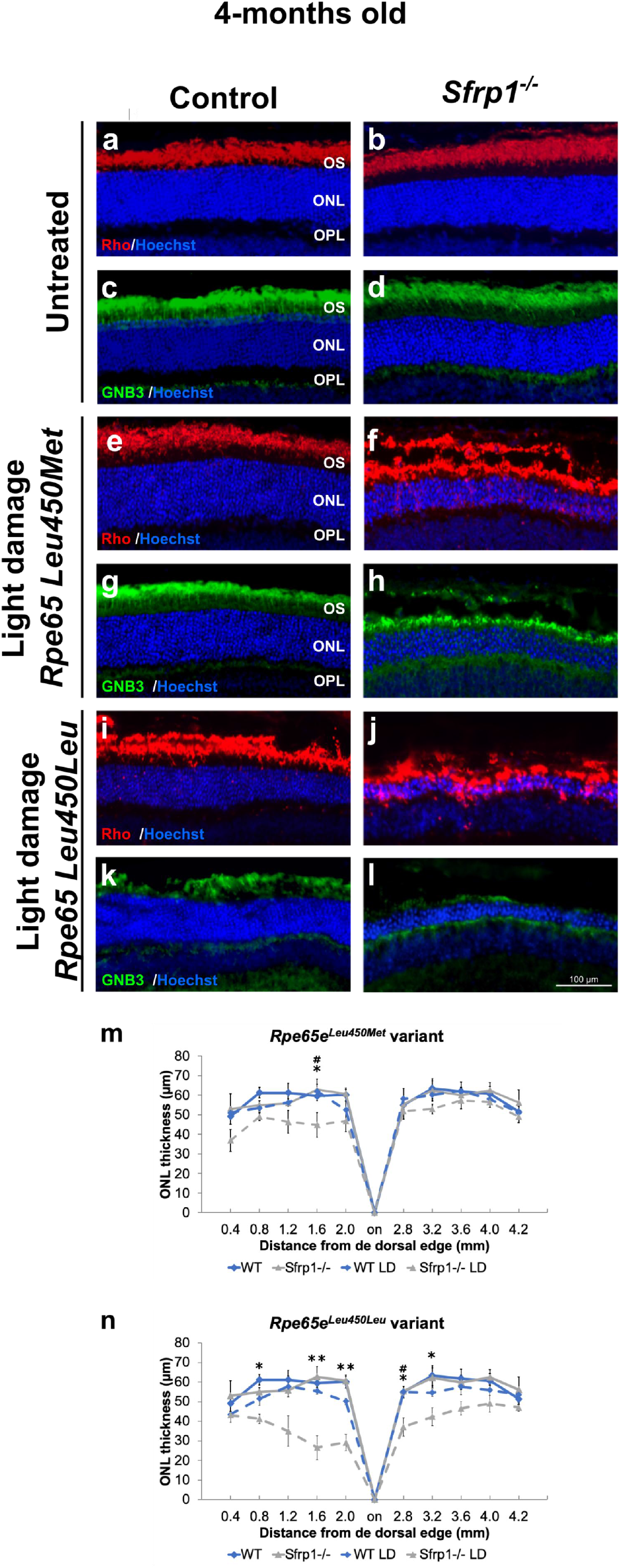
*Sfrp1^-/-^* retinas are more prone to light-induced damage even in the presence of the *Rpe65^Leu450Met^* protective variant. **a-l)** Frontal cryostat sections from 4 months-old control and *Sfrp1^-/-^* retinas carrying the *Rpe65^Leu450Leu^* or the protective *Rpe65e^Leu450Met^* variants before and 7 days after light induced-damage (light conditions: 15000 lux of cool white fluorescent light for 8 hrs). Sections were immunostained with antibodies against Rho (rods, red) or GNB3 (cones, green) and counterstained with Hoechst. **m, n)** The graphs represent the ONL thickness (measures were taken in cryostat frontal sections of the eye at the level of the optic disk) in the different conditions. Kruskal Wallis with post hoc Dunn. *indicates significant differences between *Sfrp1^-/-^* untreated and light-damage mice. #indicates significant difference between *Sfrp1^-/-^* and wt animals exposed to light damage. No significant differences were found between wt untreated and light-damage mice. * p<0.05, ** p<0.01. Scale bar in l applies to all panels. Abbreviations: ONL: outer nuclear layer; OPL: outer plexiform layer; OS outer segment.

Comparative recording of ERGs responses from *Rpe65e^Leu450Leu^;Sfrp^+/+^* and *Rpe65e^Leu450Leu^;Sfrp1^-/-^* 4 months-old mice exposed or not to light damage extended this analysis. Untreated mice lacking *Sfrp1* presented a slightly reduced rod (scotopic) and cone (photopic) response to light flashes compared to *Sfrp1^+/+^* (Fig. 7a-d). Consistently the reduced loss of photoreceptors observed in *Rpe65e^Leu450Leu^;Sfrp^+/+^* after damage (Fig. 6e,c), both scotopic and photopic responses were only slightly and non-significantly reduced in these animals (Fig. 7a-d). High intensity light exposure instead strongly reduced the rod and cone response of *Rpe65e^Leu450leu^;Sfrp1^-/-^* mice (Fig. 7a-d).

**Figure 7.**
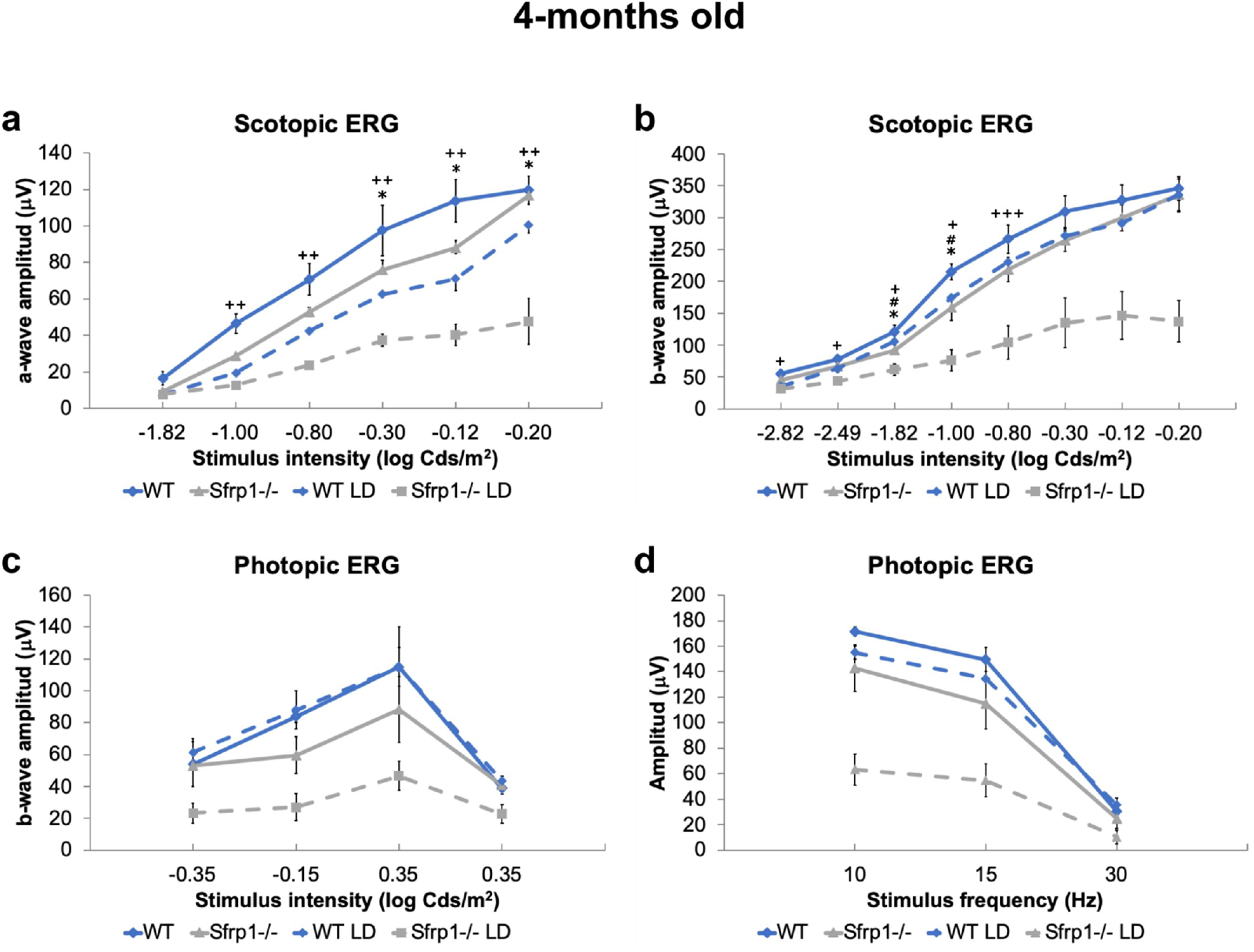
Loss of reduces the ERG response after light-induced damage even in the *Rpe65^Leu450Leu^* variant. **a-d)** The graphs show the a- and b-wave amplitude (μV) of the scotopic (rods and rod bipolar cells, a, b) and photopic (cones and cone bipolar cells, c, d) ERG response from 4 months-old WT and *Sfrp1^-/-^* animals to flash light stimuli of increasing brightness (cd/m2). Kruskal Wallis with post hoc Dunn. *indicates significant difference between *Sfrp1^-/-^* untreated and light-damage mice. #indicates significant difference between *Sfrp1^-/-^* and wt animals exposed to light damage. +indicates significant differences between *Sfrp1^-/-^* and wt untreated mice. No significant differences were found between untreated and light-damage wt mice. * p<0.05, ** p<0.01, *** p<0.001. Abbreviations: ONL: outer nuclear layer; OPL: outer plexiform layer; OS outer segment.

All in all, these data indicate that SFRP1 activity critically supports photoreceptor integrity and function upon a neurodegenerative stimulus.

## Discussion

Photoreceptors’ degeneration has devastating consequences on visual perception. So far lost photoreceptors cannot be effectively replaced but, in many cases, restraining their alterations would be sufficient to enable a normal life in affected individuals ^60^. This may be particularly important for cones because they are activated by day-light or artificial illumination and thus enable sufficient vision in current society. The identification of the mechanisms that contributes to the integrity of photoreceptors (cones in particular) may thus help designing protective/therapeutic strategies to guarantee sufficient visual percetion. In this study, we have identified SFRP1 as a component of the machinery that maintains photoreceptors’ organization and integrity, conferring them some protection against light induced damage. Our data point to a mechanism in which SFRP1 would favour the tight cellular adhesions underlying the OLM, which, in turn, is essential for photoreceptors’ function and fitness.

In absence of *Sfrp1*, cones are the first photoreceptors to show abnormalities with misplaced cell bodies and occasional OS de-compaction detected since young adult stages and light-damage susceptibility. This is somewhat unusual because cones have been shown to be more resistant to this type of damage ^57^. This unusual behaviour might reflect a particular dependence of cones on the interaction with the surround tissues ^60^. Indeed, a number of genes responsible of cone dystrophies encodes proteins involved in cell adhesion ^4, 7, 61^. Furthermore, overexpression studies in the embryonic chick retina indicate that *Sfrp1* promotes cone generation ^19^, although *Sfrp1^-/-^* eyes show no developmental defects ^18, 30, 31^. This may be because the related *Sfrp2* and *Sfrp5* compensate *Sfrp1* function during development ^12, 18, 32, 62^. In the adult mouse retina, *Sfrp2* and *Sfrp5* mRNAs are absent or poorly expressed (Fig S1), allowing to uncover a specific SFRP1 activity in the retina. We have observed that the initially subtle cone alterations worsen with time, and are followed by the appearance of rods’ defects and visual function impoverishment. The emergence of these defects have been also observed in aged retinas ^63, 64^, in which, according to our data, SFRP1 content is progressively decreasing.

SFRP proteins have been initially discovered and described as apoptosis-related proteins ^65^. We cannot fully discard that retinal SFRP1 may in part have such an activity. However, whereas SFRP2 and SFRP5 have anti-apoptotic functions ^65, 66^, SFRP1 overexpression increases cell sensitivity to pro-apoptotic stimuli ^65^, making the defects observed in *Sfrp1^-/-^* retina hardly interpretable through an apoptotic mechanism. Inhibition of Wnt signalling, a SFRP1 function observed in many different contexts ^11^, is also unlikely in the adult retina, because photoreceptors are significantly less vulnerable to light-induced damage upon Wnt signalling activation ^67^, which is the opposite of what we observe in *Sfrp1^-/-^* mice. The *Sfrp1^-/-^* retinal phenotype can instead be best explained considering the inhibitory function that SFRP1 exerts on the metalloprotease ADAM10 ^12, 13^. In a speculative model (Fig. 8), we envisage that SFRP1, produced by photoreceptors and possibly Müller glial cells, down-regulates ADAM10 activity at the OLM, thereby maintaining the required tight adhesion between the end-feet of Müller cells and the IS of the photoreceptors. In its absence, the increased proteolysis of the OLM and OS components loosen the OLM, progressively impairing photoreceptors’ anchorage. The OLM confers mechanical strength to the retina and limits the diffusion of components of the photo-transduction cascade. Its loosening sets the conditions for loss of photoreceptors’ fitness, making the cells more sensitive to light, and possibly, other types of stress (Fig. 8). This tentative model is supported by the appropriate localization of the required molecular machinery -ADAM10, SFRP1 and different cell adhesion molecules-within the retina ^44, 45, 53, 54^. Indeed, our data shows that both proteins localize in the OLM and regions of the INL compatible with Müller glial cell localization. We have previously shown that SFRP1 physically interacts with ADAM10 in both the retina and the telencephalon ^12, 13^, down-modulating the proteolysis of different ADAM10 substrates ^12, 13, 22, 24^. Consistent with this notion, here we have observed an increased shedding of N-cadherin and PCDH21 in the *Sfrp1^-/-^* retinas. Increased proteolysis of these molecules seems to weaken cell-cell interactions ^50, 51^ and thus may well explain the intermittent breakage of the OLM in the *Sfrp1^-/-^* retinas, with the consequent disorganization of the ONL and rhodopsin distribution. The latter defect is considered both a consequence and a further cause of rod degenerative signs, including the decompaction of the OS disc membrane stacks ^68^, as we observed in the *Sfrp1^-/-^* mutants.

**Figure 8.**
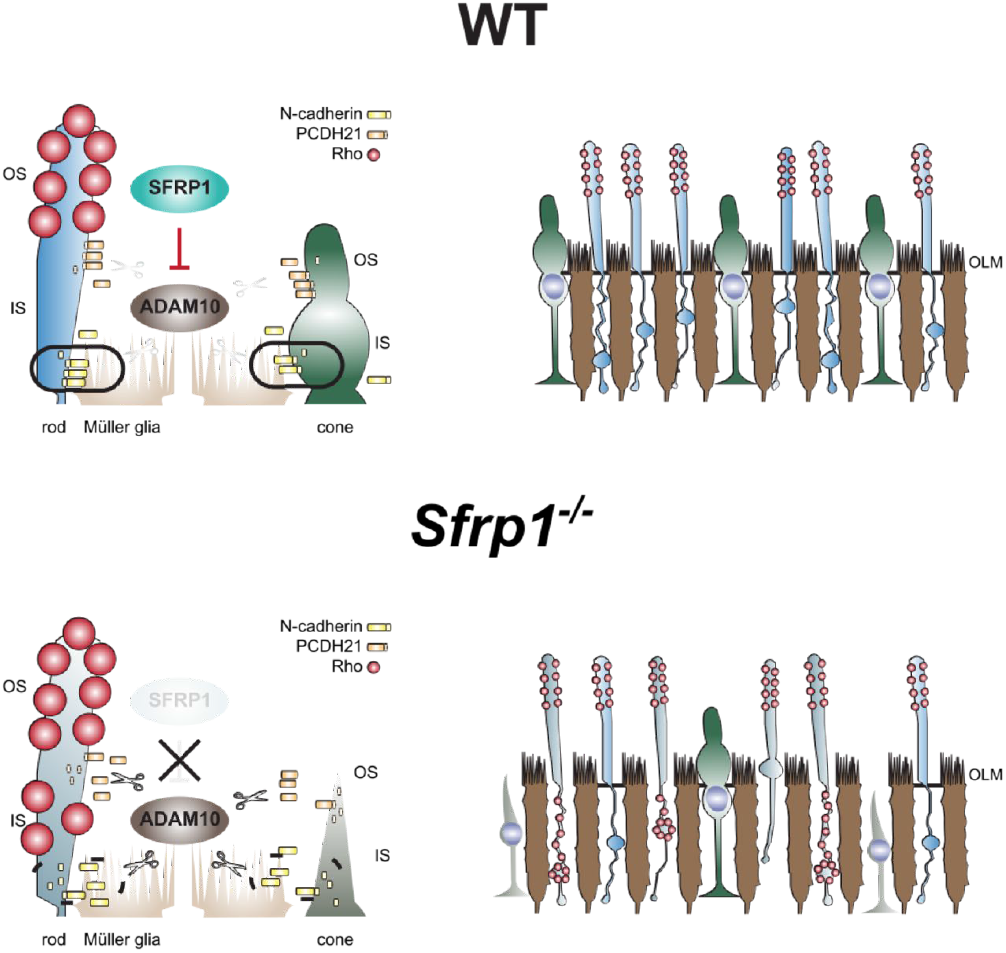
Proposed model for the defects observed in the *Sfrp1^-/-^* mice. See the discussion for details. Abbreviations: IN: inner segment; OLM: outer limiting membrane; OS: outer segment. (Author ‘s drawing)

In further support for our findings, missense mutations in PCDH21 have been identified in patients affected by autosomal recessive cone-rod dystrophy ^4, 61^, suggesting that this protocadherin might be particularly relevant for cone survival. Indeed, its targeted disruption in mice causes OS fragmentation followed by a relatively mild decrease of the ERG response and photoreceptors’ loss ^52^, somewhat resembling the *Sfrp1^-/-^* retinal phenotype. However, the precise relationship between PCDH21 shedding and photoreceptor degeneration might not be so straightforward. Analysis of PCDH21 shedding in mice that lack peripherin (*rds^-/-^*), a structural OS protein, revealed higher levels of protein expression and a decreased proteolysis ^52^ and we show that in *rd10* mice PCDH21 proteolysis was similar to wt. The reasons for these differences likely reflect the primary genetic causes of photoreceptors degeneration in *rd10* and *rds* mice: PCDH21 and peripherin have structural functions whereas *rd10* mice carry a mutation in a component of the rod phototransduction cascade ^55^. Changes in cell adhesion might thus be a consequence rather a cause in *rd10* mice.

Our study highlights a specific role of *Sfrp1* in the adult retina. We have interpreted these findings in light of the idea that SFRP1 acts as an endogenous inhibitor of ADAM10 in other contexts ^12, 13, 22, 24^. Our findings however are in apparent contradiction with the observation that in the brain SFRP1 expression increases with aging ^69^ and is upregulated in individual affected by Alzheimer’s disease ^13^, whereas it is barely detectable in the cortex of young and healthy individuals. We believe that this apparent discrepancy is linked a differential distribution of the protein in homeostatic conditions and the drastic alterations that occur during neurodegenerative events. In Alzheimer’s disease, activated glial cells (particularly astrocytes) produce SFRP1, leading to increased levels of toxic products of APP processing ^13^, which further sustain chronic inflammation ^16^. In the adult retina, both neurons and likely Müller glial cells produce SFRP1, which is localized at the OLM with a putative homeostatic function over ADAM10. Its loss of function leads to subtle and slow alterations that are not associated with glial cell reactivity. Drastic photoreceptors’ degeneration, leads instead to neuroinflammation ^70–72^. We expect that in this condition *Sfrp1* expression would be also up-regulated in reactive retinal glia, explaining the reported elevated expression and ectopic distribution of SFRP1 in RP ^25, 27^. We also expect that neuroinflammatory functions of SFRP1 ^16^ would override its homeostatic role, although testing whether this is the case is not straightforward. *SFRP1* expression is under strong epigenetic regulation ^28^, which may further tune SFRP1 protective or offensive role in the retina.

In conclusion, we show that Sfrp1 contribute to adult retinal homeostasis maintaining the integrity of the OLM. The phenotype observed in *Sfrp1* null mice suggest the possibility that the age-related decrease of retinal fitness and visual performance might involve a progressive down-regulation of *Sfrp1* expression. Furthermore, differential *SFRP1* expression among individuals may represent one of the many possible causes of the low genotypephenotype correlation observed among individuals suffering of retinal dystrophies.

## Methods

### Animals

*Sfrp1^tm1Aksh^* (*Sfrp1^-/-^* mice, thereafter) and *Sfrp5^tm1Aksh^* mice were generated as described ^30, 73^}, were kindly provided by Dr. A. Shimono (TransGenic Inc. Chuo, Kobe, Japan) and maintained in a mixed C57BL/6 x 129 genetic background (B6;129). Mutant mice were compared with wild type (wt) littermates. All animals were housed and bred at the CBMSO animal facility in a 12 h light/dark cycle and treated according to European Communities Council Directive of 24 November 1986 (86/609/EEC) regulating animal research. All procedures were approved by the Bioethics Subcommittee of Consejo Superior de Investigaciones Científicas (CSIC, Madrid, Spain) and the Comunidad de Madrid under the following protocol approval number (PROEX 100/15; RD 53/2013).

### Genotyping for Rpe65 and Crb1 variants

To determine which are the *Crb1* and *Rpe65* gene variants present in the genome of the mouse line used in this study, tail genomic DNA was isolated from *Sfrp1^-/-^* mice and their wt littermates according to standard protocols and genotyped as described ^36, 59^. To determine the sequence of *Crb1* and *Rpe65* we used the following primer pairs: *Crb1* Fw, 5’-GGTGACCAATCTGTTGACAATCC-3’; Rv, 5’-GCCCCATTTG CACACTGATGAC-3’ ^36^; *Rpe65;* Fw, 5’-CTGACAAGCTCTGTAAG-3’; Rv, 5’-CATTACCATCAT CTTCTTCCA-3’ ^36, 59^. Amplified fragments were sequenced (Secugen S.L., Madrid) and analysed using CLC Sequence Viewer 6 (Qiagen).

### In situ hybridization (ISH) and immunohistochemistry (IH)

Mice were anesthetized with sodium pentobarbital and perfused intracardially with 4% PFA in 0.1 M phosphate buffer (PB). Alternatively, animals were euthanized by CO_2_ inhalation and eyes were enucleated and fixed by immersion in 4% PFA in PB for 2 h. The anterior segment of the eye was removed and the eyecups were cryoprotected by equilibration in 15-30% sucrose gradation in PB, embedded in 7.5% gelatine/15% sucrose solution, frozen and sectioned at 16 μm. Sections were processed for in situ hybridization (ISH) as described ^22^ using a DIG-labelled anti-sense riboprobe for *Sfrp1, Sfrp2* and *Adam10*. When processed for immunohistochemistry (IH), the sections were incubated for 2 h at room temperature (RT) in a solution containing 1% BSA, 0.1% gelatine and 0.1% Triton X-100 in phosphate buffer saline (PBS). The sections were then incubated with primary antibodies diluted in blocking solution at 4°C overnight. Primary antibodies were the following: rabbit anti-SFRP1(1:500), rabbit anti-GFP (Molecular Probes, 1:1000); rabbit anti-ADAM10 (1:500, Abcam); mouse anti-Rhodopsin K16-155C ^74^ (kindly provided by Dr. P. Hargrave, 1:100), rabbit anti-Guanine Nucleotide Binding protein (G protein Beta 3 subunit, GNB3; Abcam; 1:1000), rabbit anti-β-catenin (Abcam, 1:1000 mouse anti-N-cadherin (Invitrogen; 1:500), mouse anti-Glutamine Synthetase (GS; Millipore, 1:1000). Sections were washed and incubated with the appropriate Alexa-Fluor 488 or 594-conjugated secondary antibodies (Molecular Probes, 1:1000) at RT for 90 min. After washing, the tissue was counterstained with Hoechst (Sigma), mounted in Mowiol and analysed with light (Leica DM500) or confocal microscopy (Zeiss LSM 710). Confocal images were acquired with identical settings. Figures display representative images. Every staining was performed in a minimum of 3 control and 3 mutant mice, obtaining equivalent results. Retina thickness was measured using Hoechst stained cryostat sections at 400 to 1800 μm from the optic nerve head at intervals of 200 μm. Mean ± SEM thickness calculated and plotted using Excel software. Student’s t-test was applied for comparison between wt and mutant animals.

### ELISA

To quantify the amount of SFRP1 protein, retinas from WT mice were isolated and lysed in RIPA buffer. SFRP1 protein levels were determined using a specific capture ELISA as detailed in ^13^.

### Transmission electron microscopy

Eyes were fixed by immersion in 2% glutaraldehyde and 2% PFA in 0.1 M cacodylate buffer. Samples were treated with 1% osmium tetroxide, dehydrated and embedded in Epon 812. Semi-thin sections (0.7 μm) were stained with toluidine blue and observed by light microscopy (Leica DM500), whereas ultrathin sections were analysed by electron microscopy (Zeiss EM900). Distributions of cone nuclei were measured in central retina semi-thin sections of 3 wt or mutant mice at (2 or 6 months old) and statistics was calculated and plotted using Excel software. Student’s t-test was applied for comparison between wt and mutant animals. At least 3 wt or mutant mice were analysed using electron microscopy. Figures display representative images.

### Western blot analysis

Isolated retinal tissue including the RPE was incubated in 2 mM CaCl_2_ at room temperature for 30 min and then in buffer containing 150 mM NaCl, 50 mM Tris pH 7.5, 1 mM EDTA, 1% NP-40, complete protease inhibitor cocktail and PMSF (Roche), as previously described ^52^. Total proteins (50 μg) were separated by SDS-PAGE and transferred onto PVDF membrane. Membranes were immersed in a solution of 5% non-fat milk and 0.1% Tween 20 in Tris Buffered Saline (TBS) for 2 h and then incubated with the appropriate primary antibodies diluted in blocking solution at 4°C overnight. The primary antibodies used were: rabbit antiserum against the C-terminal domain of PCDH21 (a kind gift of Dr. A. Rattner; 1:10.000), mouse monoclonal antibody generated against the C-terminal domain of N-cadherin (Invitrogen; 1:500); anti-CRB1 (a kind gift of Dr. J. Wijnholds, 1:500), and mouse anti-(α-tubulin (Sigma, 1:10.000), used as a loading control. The membranes were washed and incubated with horseradish peroxidase-conjugated rabbit or mouse antibodies followed by ECL Advanced Western Blotting Detection Kit (Amersham). The immune-reactive bands were quantified by densitometry and the amount of proteolyzed fragment was estimated as the ratio of C-terminal fragment (CTF)/full length (FL)/α-tubulin band values. Western blots were repeated at least 3 times obtaining similar results.

### Light-induced retinal damage

The pupils from 4 and 9 months-old mice were dilated by topical administration of 1% tropicamide (Alcón cusí) 30 min before light exposure and by a second administration after the first 4 h of exposure. The light exposure device consisted of cages with reflective interiors, in which freely moving animals were exposed, at the same time of the day, to 15000 lux of cool white fluorescent light for 8 h. Control animals were sacrificed before light exposure, whereas experimental animals were sacrificed 7 days after completed light exposure.

### Electroretinography (ERG)

Mice were dark-adapted overnight. All procedures were performed under dim red light following the ISCEV guidelines ^75^. Animals were anesthetized by intra-peritoneal injection of ketamine (95 mg/Kg) and xylazine (5 mg/Kg). The body temperature was maintained with an electric heating pad. The pupils were dilated by topical administration of 1% tropicamide as above. A drop of 2% methylcellulose was then placed between the cornea and the silver electrode to maintain conductivity. VisioSystem Veterinary set up (SIEM Bio-Médicale) was used to deliver flash stimuli, as well as to amplify, filter and analyse photoreceptor responses. Rod responses (Scotopic ERG) were determined using stimulation intensities ranging from −2.8 to −0.2 log Cd s/m^2^. Cone responses (Photopic ERG) were registered after 10 min adaptation to a 3 Cd/m^2^ background light using stimulation intensities ranging from −0.35 to 0.35 log Cd s/m^2^ and after 10 min adaptation to a 30.3 Cd/m^2^ background light using a stimulation intensity of 0.35 log Cd s/m^2^. Rods may respond even in photopic conditions ^76, 77^. We thus tested cone response also by flicker ERG using different frequencies that allow discriminating cone from rod response. Indeed, rods only respond to low frequency stimuli (5-15 Hz) whereas cones can respond to both low and high frequencies (5-35 Hz)^76, 77^. Flicker ERG was performed with a stimulation intensity of 0.5 log Cd s/m^2^ at frequencies of 10, 20 and 30 Hz, after 10 min adaptation to a 30.3 Cd/m^2^ background light. ERGs were recorded in 4 months old mice, 5 wt and 6 mutant mice homozygotes for the protective *Rpe65^Leu450Met^* gene variant, in 5 wt and 5 mutant mice heterozygotes for the protective *Rpe65^Leu450Met^* gene variant and in 2 wt and 3 mutant mice without protective gene variant. ERGs were also recorded in 9 months old mice, 5 wt and 5 mutant mice homozygotes for the protective *Rpe65^Leu450Met^* gene variant. Mean ± SEM values were calculated and plotted using Excel software. Mann-Whitney U test was applied for comparison between wt and mutant mice whereas Kruskal Wallis with post hoc Dunn was applied in the case of light-damaged experiments.

## Declarations

### Ethics approval

Animals were housed and treated according to European Communities Council Directive of 24 November 1986 (86/609/EEC) regulating animal research. All procedures were approved by the Bioethics Subcommittee of Consejo Superior de Investigaciones Científicas (CSIC, Madrid, Spain) and the Comunidad de Madrid under the following protocol approval number (PROEX 100/15; RD 53/2013).

### Consent for publication

Not applicable

### Availability of data and materials

All data generated or analysed during this study are included in this published article

### Authors’ contributions

EC and FM designed the experiments, performed and analysed experiments and contributed to manuscript preparation. JRC participated in immune-histochemical and ERG studies. CL performed and analysed EM studies. GP, AV and RS performed ELISA studies. MJB and AS genotyped animals and performed biochemical analysis. PE participated in experimental design, result discussion and manuscript preparation. PB conceived the study, contributed to experimental design and interpretation and wrote the manuscript.

## Acknowledgements

The authors thank O. Herreras (Cajal Institute, CSIC) and E. Martín (Universidad de Albacete) for advice and support in the initial ERG studies.

