## Supplementary material for "Sfrp1 deficiency makes retinal photoreceptors prone to degeneration": Supple Files

Supplementary Information

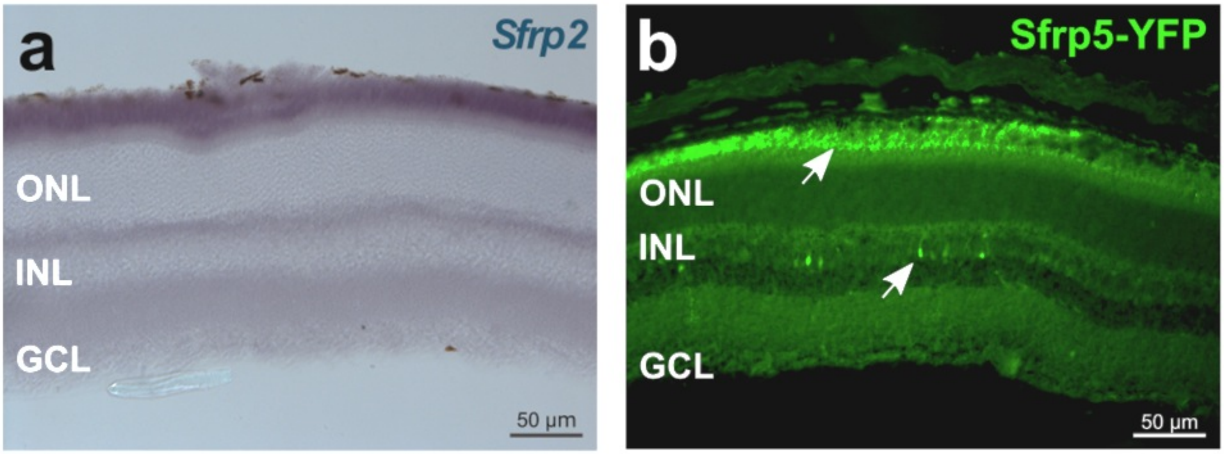


**Figure S1. Extended information from Figure 1. a)** Frontal cryostat section from 1 month-old wt animal hybridized with a probe for *Sfrp2*. No specific signal was observed. b) Frontal cryostat section from 1 month-old *Sfrp5*$^{tm1Aksh}$ animal immunostained for YFP reporter distribution. Note the presence of fluorescent signal in the RPE and sparse cells of the INL.

a

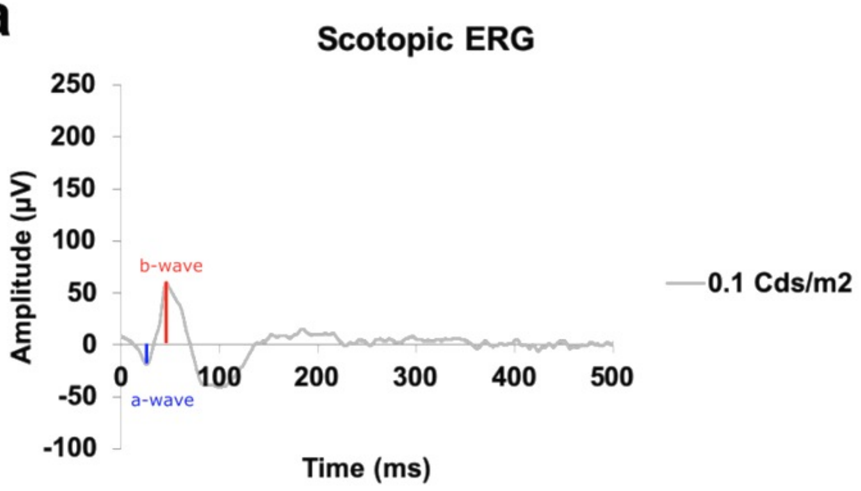


**Figure S2. Extended information from Figure 2. Typical recording under scotopic conditions were a (blue) and b wave amplitudes (red) have been indicated.**

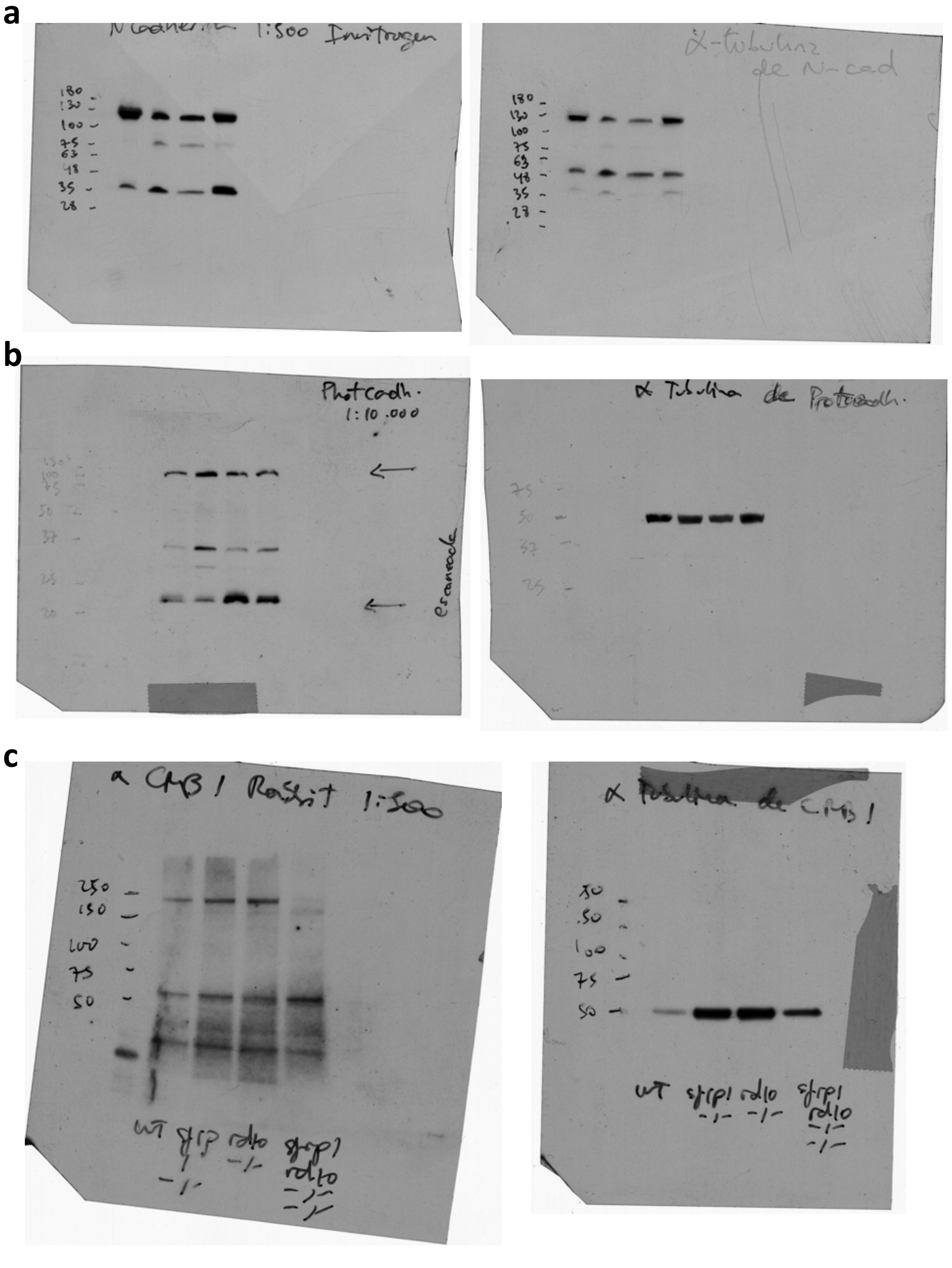


**Figure S3. Extended information from Figure 5. a-c)** Uncropped Western Blot images displayed in Fig. 5c, d, e (left) respectively together with their relative controls (right).
